# Engineering a super-secreting strain of *Escherichia coli* by directed co-evolution of the multiprotein Tat translocation machinery

**DOI:** 10.1101/2021.04.25.441317

**Authors:** May N. Taw, Mingji Li, Daniel Kim, Mark A. Rocco, Dujduan Waraho-Zhmayev, Matthew P. DeLisa

## Abstract

*Escherichia coli* remains one of the preferred hosts for biotechnological protein production due to its robust growth in culture and ease of genetic manipulation. It is often desirable to export recombinant proteins into the periplasmic space for reasons related to proper disulfide bond formation, prevention of aggregation and proteolytic degradation, and ease of purification. One such system for expressing heterologous secreted proteins is the twin-arginine translocation (Tat) pathway, which has the unique advantage of delivering correctly folded proteins into the periplasm. However, transit times for proteins through the Tat translocase, comprised of the TatABC proteins, are much longer than for passage through the SecYEG pore, the translocase associated with the more widely utilized Sec pathway. To date, a high protein flux through the Tat pathway has yet to be demonstrated. To address this shortcoming, we employed a directed co-evolution strategy to isolate mutant Tat translocases for their ability to deliver higher quantities of heterologous proteins into the periplasm. Three super-secreting translocases were selected that each exported a panel of recombinant proteins at levels that were significantly greater than that observed for wildtype TatABC or SecYEG translocases. Interestingly, all three of the evolved Tat translocases exhibited quality control suppression, suggesting that increased translocation flux was gained by relaxation of substrate proofreading. Overall, our discovery of highly efficient translocase variants paves the way for the use of the Tat system as a powerful complement to the Sec pathway for secreted production of both commodity and high value-added proteins.

## Introduction

The export of recombinant proteins out of the cytoplasm and into the periplasm of Gram-negative bacteria is a commonly used strategy in preparative protein expression ^1-3^ and in numerous protein engineering applications ^4-13^. In these and many other instances, the general secretory (Sec) pathway has served as the primary route by which heterologous proteins are delivered to the periplasmic space. Sec export involves translocation through a narrow (5–8 Å) diameter pore formed by the SecYEG translocase, which necessitates that protein substrates attain an unfolded conformation prior to translocation and be capable of reaching their native, biologically active conformation after they have been ‘threaded’ through the Sec pore ^14, 15^. These requirements render many Sec-targeted recombinant proteins incompetent for export, either because they fold too quickly or tightly in the cytoplasm or become stalled in the secretion apparatus ^16-19^. Still others transit the Sec machinery only to undergo extensive degradation or misfolding in the periplasmic compartment ^20, 21^. Hence, only a subset of all proteins are compatible with the Sec pathway, which serves to limit the functional secretory capacity of *E. coli* cells.

The twin-arginine translocation (Tat) pathway sits alongside the Sec system in the bacterial cytoplasmic membrane and provides an alternative route by which native and heterologous proteins can be trafficked to the periplasm ^22-25^. Importantly, the Tat translocase, a heteromeric complex involving multiple copies of the TatA, TatB and TatC integral membrane proteins, is endowed with several unique, inbuilt characteristics that offer distinct advantages over the Sec pathway for expressing heterologous proteins. First and foremost, the Tat pathway exports fully folded substrate proteins including those that are inherently incompatible with Sec export ^26, 27^, which is in stark contrast to Sec-dependent export of extended polypeptides that have not yet reached a native conformation. In fact, proteins ranging in size from 9-150 kDa and involving two or more preassembled subunits can be accommodated by the Tat translocase ^13, 28, 29^. In addition, the Tat system has proven to be compatible with a wide range of complex biopharmaceutical products including human growth hormone (hGH), interferon (IFN)-α2b, tissue plasminogen activator (tPA), and a variety of recombinant single-chain Fv (scFv) antibody fragments ^13, 28, 30-34^. Second, a folding quality-control (QC) mechanism that discriminates between folded and mis/unfolded substrate proteins, allowing export of only the former, appears to be an inherent property of the Tat translocase ^26, 35-38^. This mechanism provides structural proofreading of substrates, quantitatively rejecting virtually all misfolded or misassembled substrates that have been tested to date ^26, 31, 36, 39-42^. Moreover, because the QC mechanism effectively blocks poorly folded polypeptides from entering the periplasm, proteins that accumulate periplasmically represent a fairly homogenous population of properly folded, highly active molecules ^43^. Hence, the Tat system provides a convenient filter that enriches product quality in a manner that should be beneficial for downstream processing.

However, despite these clear advantages, the bacterial Tat pathway remains underutilized for preparative expression of secreted proteins. One reason for this is the fact that protein flux through the *E. coli* Tat translocase is significantly lower than through the Sec translocase, with translocation rates of a few minutes for Tat substrates versus just a few seconds for Sec substrates ^39, 44, 45^. The low flux appears to be due in part to the low abundance of Tat translocases in the membrane. Indeed, expression of additional copies of the *tatABC* genes from either a plasmid or the chromosome significantly enhanced the levels of substrate export ^46, 47^, which translated to secreted titers of hGH that reached ∼5 g/L following extended fed-batch fermentation ^48^. Here, we hypothesized that low secretion flux may also be “encoded” in the Tat translocase itself, with natural designs evolving to meet the needs of the organism and not to achieve optimal biotechnological performance. If this hypothesis was correct, then it should be possible to isolate mutations in translocase components that release the brakes on translocation and lead to measurable enhancement of Tat export capacity. To test this notion, we attempted to increase secretion flux through the Tat system using a directed co-evolution strategy to engineer mutant translocases that are capable of exporting greater amounts of heterologous proteins into the periplasm. Three super-secreting translocases were selected that each exported significantly greater quantities of a human scFv antibody, human growth hormone, and several other recombinant proteins compared to export of the same proteins by wildtype (wt) translocases. Interestingly, increased translocation flux for all three evolved translocases was accompanied by relaxation of substrate proofreading, an outcome reminiscent of enzyme evolution dynamics whereby adaptation towards a new function involves a trade-off with the original function ^49^. We anticipate that Tat translocase variants that secrete exceptional amounts of recombinant proteins into the periplasm will prove to be useful for a wide range of applications ranging from laboratory and preparative protein production to high-throughput screening of combinatorial libraries, all of which hinge critically on efficient protein export.

## Results

### Genetic selection of super-secreting Tat translocases

To engineer the Tat translocation machinery for enhanced export of proteins, we devised a directed co-evolution approach that focused on the TatABC proteins because these are the minimal components required for bilayer translocation of substrate proteins. At the heart of our approach was a bacterial genetic selection that linked Tat-dependent export of scFv antibodies with antibiotic resistance ^33^, which we hypothesized here would enable facile isolation of mutant translocases simply by demanding bacterial growth on plates supplemented with antibiotic. This genetic selection leveraged plasmid pSALect-scFv13-R4, which encoded a chimeric reporter comprised of a single-chain Fv antibody specific for β-galactosidase (β-gal), called scFv13-R4, modified at its N-terminus with the Tat-dependent signal peptide from *E. coli* TorA (spTorA) and at its C-terminus with the selectable marker enzyme TEM1 β-lactamase (Bla) (**Fig. 1a**) ^33^. It should be noted that scFv13-R4 was evolved for efficient cytoplasmic folding in the absence of disulfide bonds ^50^ and, as a result, can be efficiently exported from the cytoplasm of wildtype *E. coli* ^33^. When the resulting spTorA-scFv13-R4-Bla chimera was expressed in *E. coli* DADE cells ^51^ in which all of the *tat* genes were deleted from the genome and instead supplied on a pBR322-based plasmid, a low level of resistance was observed following spot-plating of cells on Luria-Bertani (LB) agar supplemented with a range of carbenicillin (Carb) concentrations (**Supplementary Fig. 1**). Despite the plasmid-based overexpression of the Tat machinery, the resistance of these cells was still below that conferred by a Sec-targeted version of the same substrate (spDsbA-scFv13-R4-Bla) expressed in DADE cells with physiological levels of SecYEG (**Supplementary Fig. 1**), confirming the markedly lower export efficiency of Tat relative to Sec.

**Figure 1.**
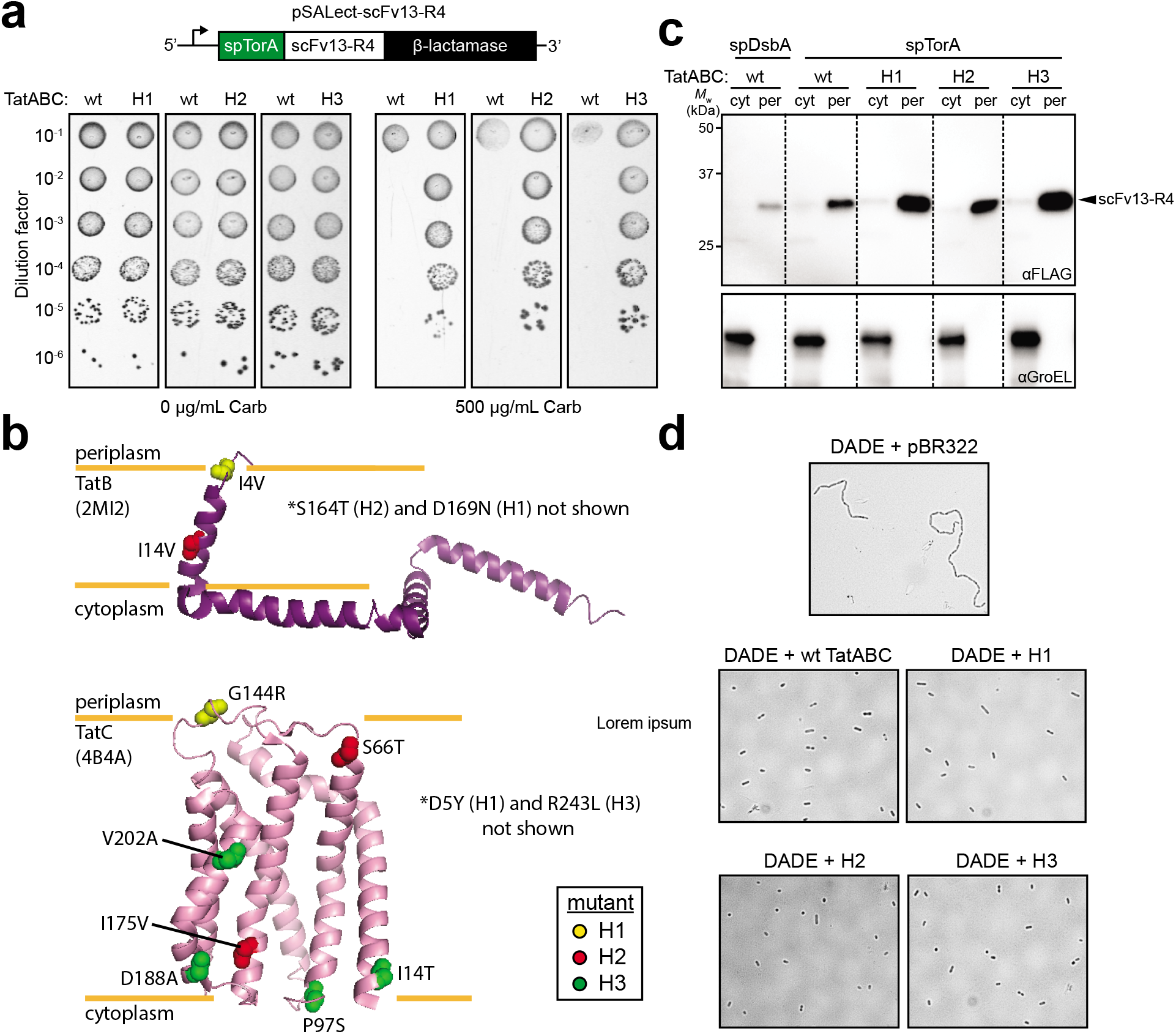
Genetic selection of super-secreting TatABC translocase mutants. (a) Schematic of spTorA-scFv13-R4 chimera cloned in plasmid pSALect ^52^. Spot plating of DADE cells co-expressing spTorA-scFv13-R4-Bla along with either wt TatABC or one of the isolated translocase mutants as indicated. Serially diluted cells were spotted on LB-agar plates supplemented with 500 μg/mL carbenicillin (Carb) or 20 μg/mL tetracycline and 30 μg/mL chloramphenicol (0 μg/mL Carb). (b) Ribbon diagram of the solution structure of *E. coli* TatB adapted from Zhang *et al*. ^53^ and the *Aquifex aeolicus* TatC crystal structure from Rollauer *et al*.^54^ Mutations isolated for enhanced export in mutants H1, H2, and H3 are marked by yellow, red, and green spheres, respectively. A few mutations not represented in the structures are denoted by an asterisk. (c) Western blot analysis of cytoplasmic (cyt) or periplasmic (per) fractions prepared from DADE cells co-expressing the FLAG-6xHis-tagged version of spTorA-scFv13-R4 along with either wt TatABC or one of the isolated translocase mutants as indicated. Fractions corresponding to an equivalent number of cells were loaded in each lane. Anti-FLAG antibody was used to detect scFv13-R4, while anti-GroEL antibody was used to confirm an equivalent loading of samples in cytoplasmic lanes and that cytoplasmic contents did not significantly contaminate the periplasmic fractions. Molecular weight (*M*_W_) marker is indicated on the left. (d) Light microscopy of DADE cells carrying empty pBR322 or a plasmid encoding either wt TatABC, H1, H2, or H3 translocases as indicated. All results are representative of at least three biological replicates.

Towards our goal of isolating more efficient Tat translocases, we hypothesized that cells expressing TatABC variants with the ability to translocate greater amounts of spTorA-scFv13-R4-Bla would be significantly more resistant to Carb compared to cells co-expressing the same reporter in the presence of wild-type (wt) TatABC due to increased levels of the Bla fusion partner in the periplasm of the former. To test this hypothesis, DADE cells were co-transformed with pSALect-scFv13-R4 along with a plasmid library of *tatABC* genes that was generated by cloning an error-prone PCR library of the *tatABC* operon into plasmid pBR322 ^36^. Selection of super-secreting TatABC variants was performed by plating the resulting cell library, containing ∼5.5×10^6^ members, on LB agar plates supplemented with Carb at a concentration (500 μg/mL) that was observed to inhibit growth (**Supplementary Fig. 1**). To remove false positives that may have arisen due to host mutations, all colonies that grew under selection conditions were cured of plasmid pSALect-scFv13-R4 so that the remaining translocase plasmid could be directly isolated. Following isolation, each pBR322-based translocase plasmid was used to freshly transform DADE cells carrying pSALect-scFv13-R4 and the resulting resistance phenotypes were confirmed through spot plating. Following this procedure, three unique translocase mutants named H1, H2, and H3 were isolated, all of which conferred significantly greater Carb resistance to DADE cells expressing spTorA-scFv13-R4 (**Fig. 1a**). In all cases, the resistance of cells grown on plates lacking Carb was nearly identical, indicating that the different Carb-resistant phenotypes were specific to Tat-mediated export of scFv13-R4 and not reduction in fitness caused by its expression. Sequencing of the complete *tatABC* operons corresponding to these positive clones revealed that clones H1 and H2 harbored missense mutations in the *tatB* and *tatC* genes while clone H3 carried mutations only in *tatC* (**Fig. 1b**). None of the clones carried any *tatA* mutations.

To determine whether the mutations acquired by the translocase mutants affected normal Tat function, export of two native Tat substrates, the amidases AmiA and AmiC, was evaluated. These two proteins are responsible for the cleavage of peptidoglycan at the septum during cell division and are natively translocated to the periplasm via the Tat pathway ^55^. Disruption of the Tat pathway, including deletion of any of its essential components (*e*.*g*., TatA, TatB, and/or TatC), blocks AmiA/C from reaching the periplasm, which in turn causes incomplete cell division that manifests in the appearance of a chain-like phenotype when viewed by microscope ^55^. Indeed, the formation of clearly visible chains was observed for DADE cells, which are deficient in all Tat proteins (**Fig. 1d**). Importantly, when wt or mutant translocases were provided to DADE cells on plasmid pBR322, normal cell division was restored as evidenced by the appearance of only singlet and doublet cells and absence of longer cell chains (**Fig. 1d**), indicating that translocase variants were capable of properly localizing native substrates to the periplasm.

### Enhanced export by translocase variants is independent of Bla reporter

We next determined whether the superior export observed for the translocase variants was retained in the absence of the Bla reporter protein. To this end, we replaced the gene encoding the large C-terminal Bla protein in pSALect-scFv13-R4 with a dual FLAG-6xHis tag, which was introduced to facilitate detection and purification. When immunoblots of subcellular fractions were probed with an anti-FLAG antibody, significantly increased levels of scFv13-R4 were clearly observed in the periplasmic fractions isolated from cells overexpressing the mutant translocases compared to the wt TatABC machinery (**Fig. 1c**). The improved periplasmic accumulation that was triggered by the mutant Tat translocases also exceeded the amount seen in cells expressing Sec-targeted scFv13-R4 (**Fig. 1c**), indicating that the evolved translocases outperformed the native SecYEG machinery that itself was expressed at physiological levels. Collectively, these results confirmed that the H1, H2, and H3 translocase variants were each capable of greatly enhancing export of unfused scFv13-R4 to the periplasmic compartment, thereby confirming our hypothesis that the resistance phenotypes of our reporter strains were a reliable indicator of super-secreting translocases.

### Enhanced export of diverse substrate proteins

To demonstrate the practical utility of our engineered translocases, we next tested whether the enhanced export observed for scFv13-R4 could be extended to other proteins of interest (POIs). Specifically, we cloned a panel of 8 heterologous targets into plasmid pSALect (**Fig. 2a**). The panel was representative of biotechnologically-relevant POIs spanning a range of sizes (8-50 kDa) and topologies, including: (1) additional affinity binders including scFv-D10 specific for the capsid protein D (gpD) of bacteriophage lambda ^56^, designed ankyrin repeat protein (DARPin) off7 specific for *E. coli* maltose-binding protein (MBP) ^57^, and fibronectin type III (FN3) monobody YS1 also specific for MBP ^58^; (2) recombinant hGH, an FDA-approved biotherapeutic that is produced using *E. coli* ^59^ and compatible with the Tat pathway ^47, 48^; and (3) mammalian proteins or protein domains that were previously expressed via the Tat system in *E. coli* ^60^ including human CASP2 protease (hCASP2), human GATA2 transcription factor (hGATA2), the sterile alpha motif (SAM) domain of murine EphB2 (mEphB2^SAM^), and the Ras binding domain (Ras-BD) of human RAF1 (hRAF1^Ras-BD^). The resulting pSALect-based plasmids were each used to transform DADE cells carrying either wt TatABC or H2 in plasmid pBR322, and the transformants were subjected to spot plating as above. As was seen for scFv13-R4, cells expressing the 8 different POIs in the presence of the H2 translocase were significantly more Carb resistant than their wt TatABC counterparts (**Fig. 2a**), indicating broad substrate compatibility for the H2 translocase that enabled improved export for these structurally diverse protein targets. Immunoblot analysis of one of the POIs, namely spTorA-hGH, in the absence of the Bla reporter revealed clearly greater periplasmic accumulation in the presence of the H2 translocase versus wt TatABC (**Supplementary 2a**), thereby confirming that the increased resistance observed in these experiments corresponded to enhanced export of the POI. It should be noted that while hGH has two disulfide bonds, folding of the protein is remarkably similar in the presence and absence of these bonds ^61^. This helps to explain how reduced hGH, which we presume is the predominant species generated in the cytoplasm of DADE cells, is able to pass the Tat QC filter and transit the Tat pathway.

**Figure 2.**
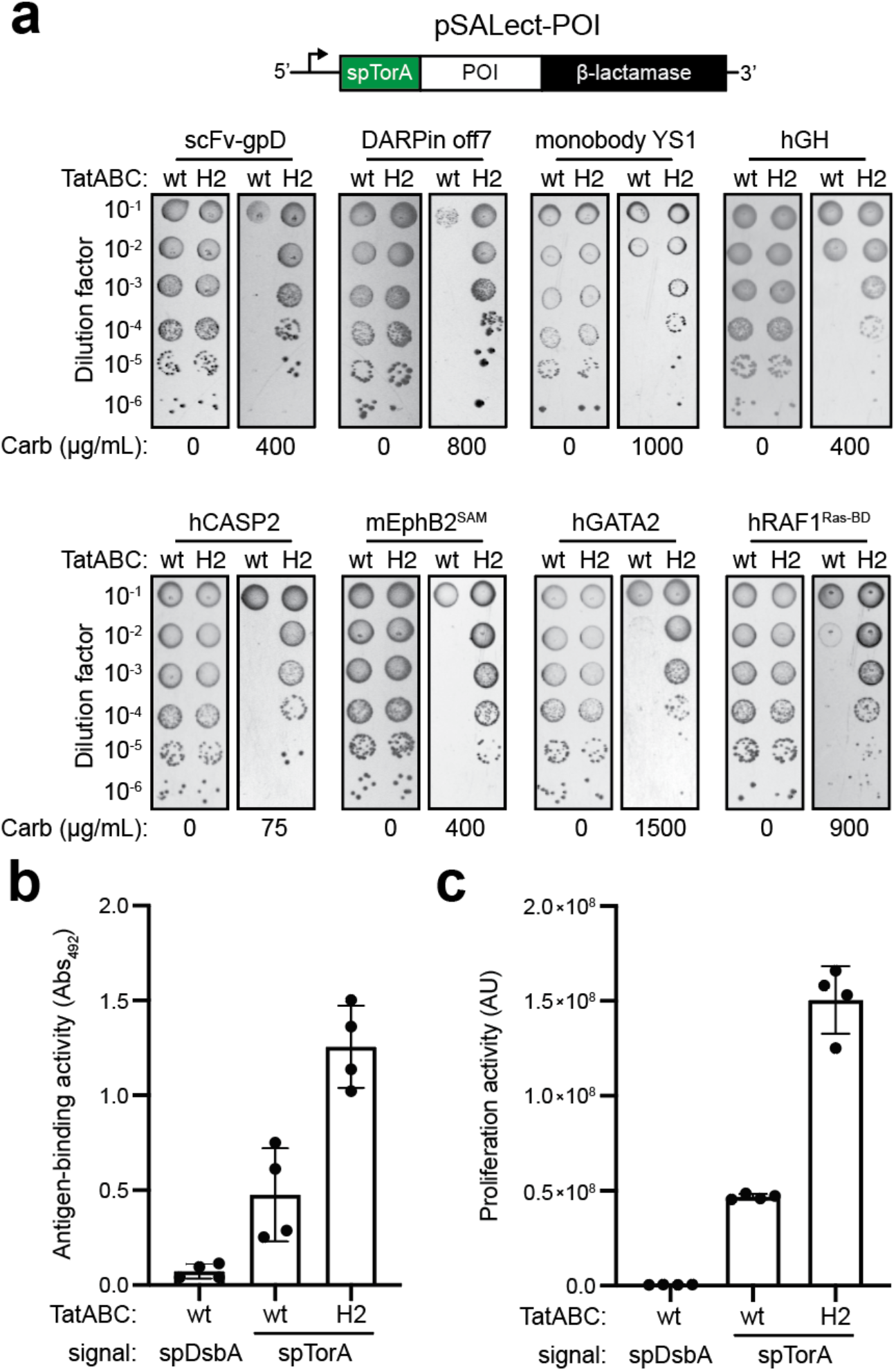
Enhanced export of scFv13-R4 and other recombinant proteins. (a) Schematic of pSALect-based constructs whereby chimeras were generated by inserting the protein of interest (POI) between spTorA and β-lactamase (Bla). Spot plating of DADE cells co-expressing different spTorA-POI-Bla chimeras along with wt TatABC or the H2 translocase as indicated. Serially diluted cells were spotted on LB-agar plates supplemented with carbenicillin (Carb) as indicated or 20 μg/mL tetracycline and 30 μg/mL chloramphenicol (0 μg/mL Carb). Results are representative of at least three biological replicates. (b) Antigen-binding activity of scFv13-R4 purified from DADE cells expressing either spDsbA-scFv13-R4 or spTorA-scFv13-R4 along with either wt or H2 translocase as indicated. Equivalent volumes of each protein were assayed for binding by ELISA with β-galactosidase (β-gal) as immobilized antigen. Data are the average of biological quadruplicates and the error bars represent the standard deviation of the mean. (c) Growth stimulatory activity of hGH purified from DADE cells expressing either spDsbA-hGH or spTorA-hGH along with either wt or H2 translocase as indicated. Ba/F3-GHR cell cultures were incubated for 72 h in the presence of equivalent volumes of hGH purified from the different strains. Data are the average of biological quadruplicates and the error bars represent the standard deviation of the mean.

The amount of secreted scFv13-R4 that was achieved as a result of the translocase variants was determined by purifying the scFv13-R4 construct from the periplasm of DADE cells carrying either the wt Tat translocase or one of the translocase variants, namely H2. Consistent with the spot plating and immunoblot results above, a total of 0.52 mg/L of Tat-targeted scFv13-R4 was recovered from the periplasm of cells carrying the H2 translocase versus only 0.07 mg/L from cells with wt TatABC (**Supplementary Fig. 3**). It should be pointed out that this latter value was in close agreement with a previously reported titer of 0.06 mg/L for Tat export of the same protein ^43^. This increase in titer corresponded to a significant difference in scFv13-R4 binding activity as determined by ELISA, with cells overexpressing H2 yielding ∼3-fold more activity compared to their wt TatABC counterparts (**Fig. 2b**). Interestingly, the titer for Sec-targeted scFv13-R4 was only 0.01 mg/L (**Supplementary Fig. 3**), which was appreciably lower than the amount recovered from cells overexpressing H2 or even wt TatABC. To determine if the enhancement seen for scFv13-R4 was generalizable to other POIs, we compared the titers of Tat-targeted hGH that was recovered from the periplasm of DADE cells carrying the different translocases. Consistent with the above results, cells carrying H2 yielded 0.48 mg/L of periplasmic hGH whereas those carrying wt TatABC produced only 0.17 mg/L (**Supplementary Fig. 2b**), which in turn corresponded to a >3-fold difference in biological activity (**Fig. 2c**). It should also be noted that relatively high purities of the Tat-targeted scFv13-R4 and hGH constructs were achieved after just a single Ni-NTA purification step, highlighting a well-known advantage of periplasmic expression that stems from the fact that there are much fewer endogenous proteins in the periplasm relative to the cytoplasm ^1, 3^.

### Super-secreting phenotype can be reduced to a single missense mutation

Each translocase variant had multiple mutations and none of the mutations overlapped, making it difficult to discern the contribution of individual mutations to the super-secreting phenotype. Therefore, to narrow the number of mutations needed for hypersecretion, we created hybrid mutant/wt translocases by systematically replacing one of the mutated *tat* alleles from each mutant with the corresponding wt alleles. For example, combining the mutant *tatB* allele from H1 along with wt *tatAC* alleles gave rise to a new translocase variant harboring only the *tatB* mutations I4V and D169N. This hybrid translocase called TatAB^H1^C was unable to recapitulate the enhanced export of scFv13-R4 as evidenced by a resistance profile upon spot plating that closely resembled wt TatABC (**Fig. 3a** and **Supplementary Fig. 4a**). Likewise, the TatABC^H1^ hybrid, which harbored only the *tatC* mutations D5Y and G144R from clone H1, was also unable to confer resistance beyond that of wt TatABC, indicating that the mutations in both TatB and TatC were essential for enhancing export. Following an identical strategy for clone H2, which also contained two mutations each in TatB (I14V and S164T) and TatC (S66T and I175V), we observed that hybrid translocases TatAB^H2^C and TatABC^H2^ both conferred resistance that was comparable to wt TatABC (**Fig. 3a** and **Supplementary Fig. 4a**). Taken together, these results suggest that the super-secreting phenotype seen for both the H1 and H2 translocases arises from a cooperative effect of the mutations in *tatB* and *tatC*.

**Figure 3.**
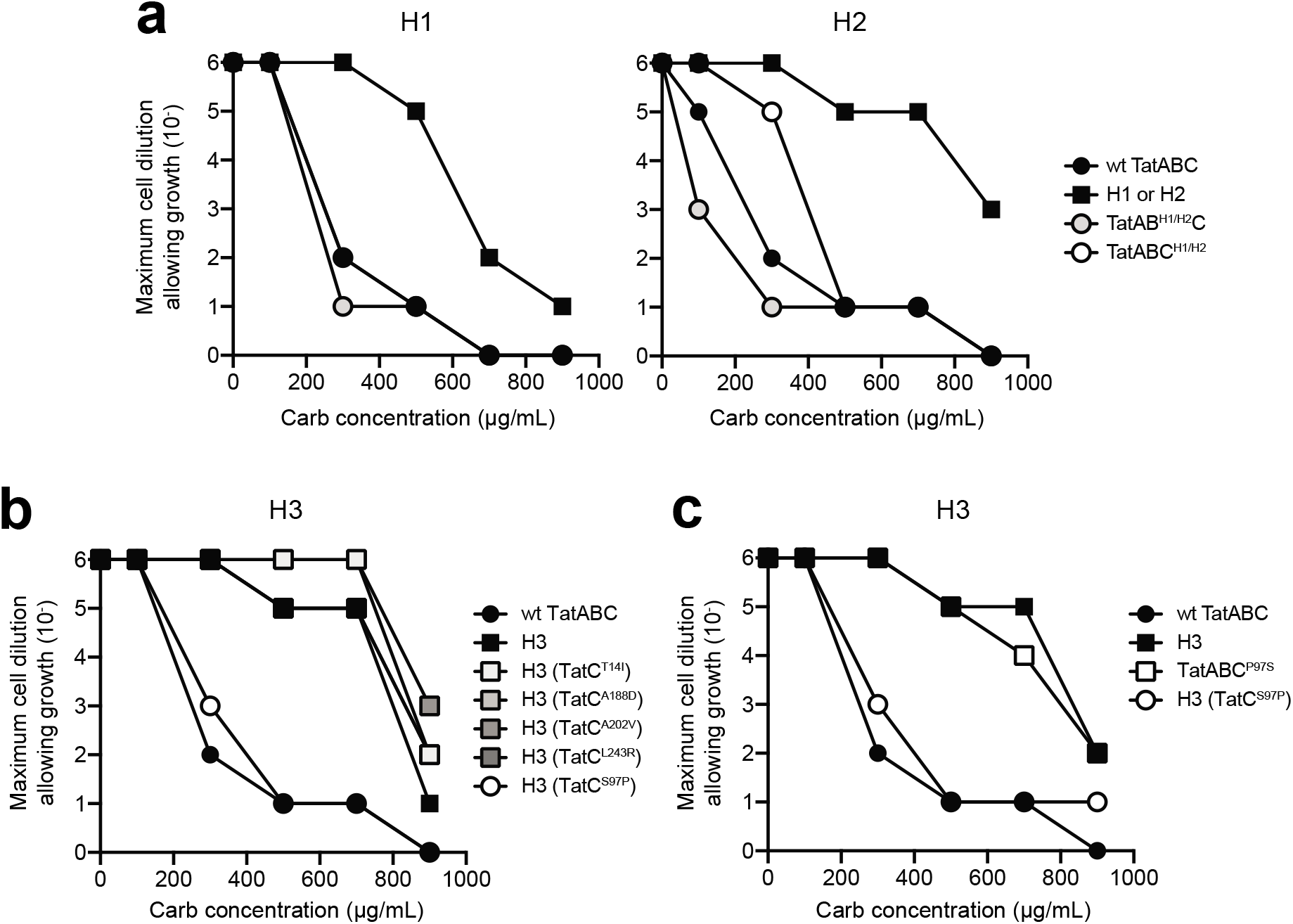
Mutations required for super-secreting phenotype of translocase mutants. (a) Spot plating of DADE cells co-expressing spTorA-scFv13-R4 along with either wt TatABC, H1, H2, or one of the hybrid translocase mutants in which the mutant *tatB* or *tatC* allele was substituted for the wt copy in the plasmid encoding wt TatABC. Kill curves were generated by spot plating serially diluted cells over a range of carbenicillin (Carb) concentrations as indicated. (b) Same as in (a) but with DADE cells co-expressing spTorA-scFv13-R4 along with either wt TatABC, H3, or one of the H3 revertant mutants as indicated. (d) Same as in (a) but with DADE cells co-expressing spTorA-scFv13-R4 along with either wt TatABC, H3, the S97P revertant of H3, or the hybrid translocase in which P97S was directly mutated in *tatC* (TatABC^P97S^). All results are representative of at least three biological replicates.

For the H3 translocase, which contained five mutations in TatC (I14T, P97S, D188A, V202A, and R243L) but none in the other Tat components, we took a slightly different approach. Specifically, each individual mutation in the *tatC* allele of H3 was reverted back to the wt amino acid by site-directed mutagenesis (*e*.*g*., P97S was mutated to S97P) and subsequently evaluated by spot plate analysis. With the exception of S97P, all of the reversion mutations retained the ability to enhance scFv13-R4 export to an extent that was indistinguishable from the H3 translocase (**Fig. 3b** and **Supplementary Fig. 4b**), indicating that these were neutral mutations with respect to the super-secretion phenotype. On the other hand, the S97P revertant exhibited a resistance profile that was comparable to wt TatABC (**Fig. 3b** and **Supplementary Fig. 4b**). To independently verify this result, we introduced the P97S mutation individually to *tatC* in the plasmid encoding wt TatABC and observed a resistance profile that was identical to that seen for the H3 translocase (**Fig. 3c** and **Supplementary Fig. 4c**), indicating that the super-secreting phenotype was entirely attributable to a single substitution at residue P97 within the second transmembrane helix of TatC.

### Translocase variants retain strict signal peptide specificity

Several of the mutations in H1, H2 and H3 including P97S were located near the N-terminus and the first cytoplasmic loop of TatC between TM2 and TM3, a highly conserved region that binds the signal peptide ^62^. Moreover, a previous search for suppressor translocases that could export substrates with defective signal peptides identified the same P97S substitution in TatC, albeit in combination with W92G ^63^. In light of this common mutation, we wondered whether enhanced export could arise through altered interactions between TatC and the signal-peptide in a manner that relaxes signal peptide specificity. To investigate this possibility, we tested translocase variants for their ability to export scFv13-R4 bearing a defective Tat signal in which the obligatory twin-arginine residues were swapped to twin lysines (spTorA^KK^-scFv13-R4). Spot plating analysis of DADE cells co-expressing spTorA^KK^-scFv13-R4 with each of the translocase variants revealed extremely weak resistance of cells to even low levels of Carb (**Fig. 4a**). These findings indicated that the H1, H2, and H3 retained specificity for authentic signal peptides and thus it appears unlikely that translocase variants operate at the level of signal peptide recognition.

**Figure 4.**
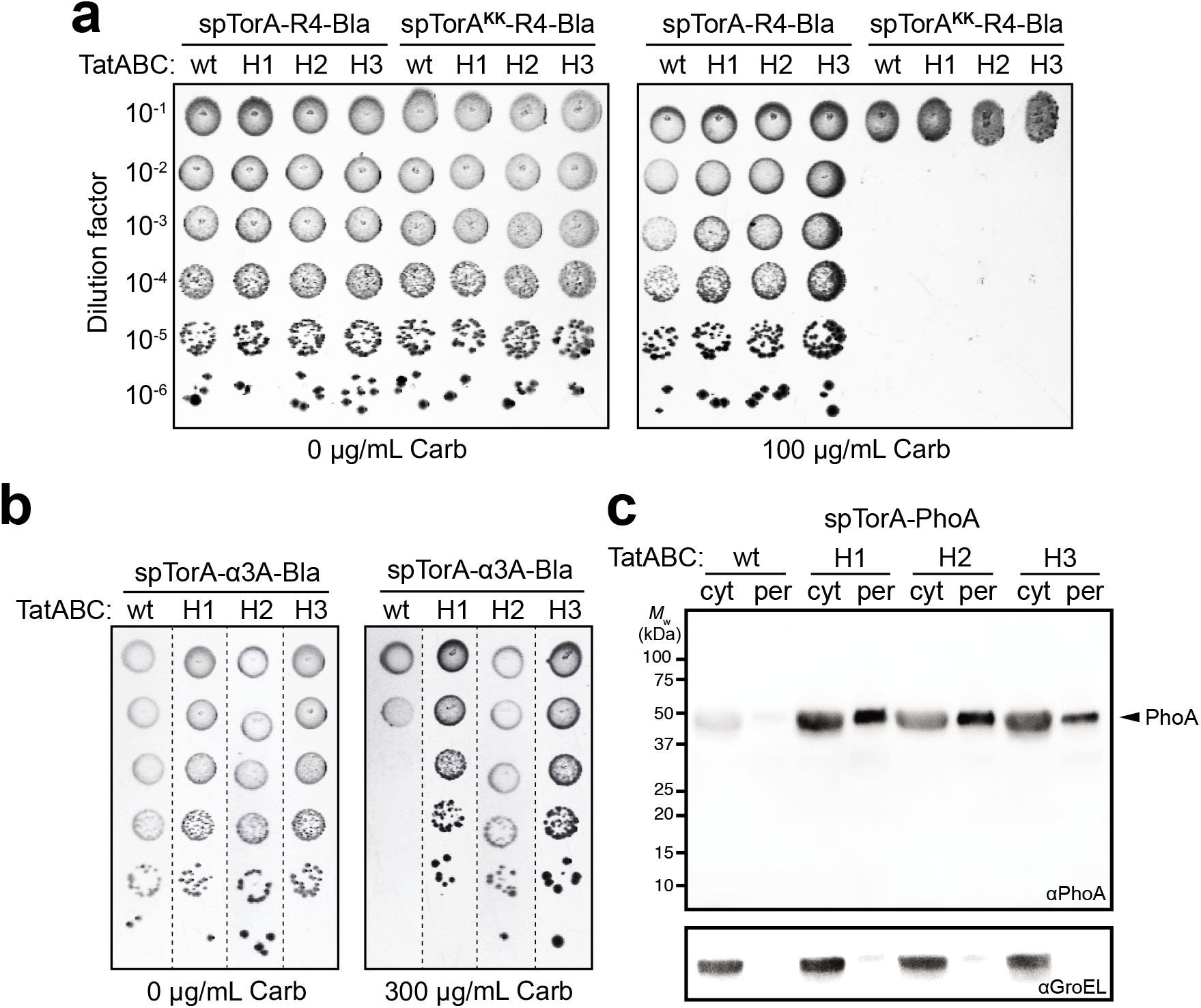
Signal peptide specificity and folding QC of translocase mutants. (a) Spot plating of DADE cells co-expressing spTorA-R4-Bla or spTorA^KK^-R4-Bla along with wt TatABC or one of the isolated translocase mutants as indicated. (b) Spot plating of DADE cells co-expressing spTorA-α_3_A-Bla along with wt TatABC or one of the isolated translocase mutants as indicated. Serially diluted cells in both (a) and (b) were spotted on LB-agar plates supplemented with 100 or 300 μg/mL carbenicillin (Carb) or 20 μg/mL tetracycline and 30 μg/mL chloramphenicol (0 μg/mL Carb). (c) Western blot analysis of cytoplasmic (cyt) and periplasmic (per) fractions prepared from DADE cells co-expressing spTorA-PhoA along with wt TatABC or one of the isolated translocase mutants as indicated. Fractions corresponding to an equivalent number of cells were loaded in each lane. PhoA was probed with anti-PhoA antibody, while anti-GroEL antibody was used to confirm an equivalent loading of samples in cytoplasmic lanes and that cytoplasmic contents did not significantly contaminate the periplasmic fractions. Molecular weight (*M*_W_) marker is indicated on the left. All results are representative of at least three biological replicates.

### Translocase variants exhibit relaxed quality control

Besides contributing to defective signal peptide suppression, the P97S mutation was also uncovered in a search for suppressor translocases capable of exporting misfolded substrate proteins, which are normally rejected by the wt translocase ^36^. The isolation of several quality control (QC) suppressors led to the conclusion that the Tat translocase sits at the center of an integrated QC system that involves “sensing” of the substrate folding state prior to membrane translocation. Given that this substrate proofreading may be a rate-limiting step in the translocation process, we speculated that inactivation of the QC mechanism could lead to faster transit times and thus greater accumulation of substrates in the periplasm, as was observed for H1, H2, and H3. To test this hypothesis, we first made use of a previously described Bla reporter construct that directly linked Tat folding QC with antibiotic resistance ^36^. This construct was comprised of a designed three-helix-bundle protein called α3A that was modified at its N-terminus with spTorA and at its C-terminus with the Bla reporter. Importantly, due to the α3A protein’s strong propensity for aggregation ^64^, this tripartite chimera serves as a reliable reporter of Tat QC. When DADE cells co-expressing spTorA-α3A-Bla and wt TatABC were subjected to spot plating analysis, we observed very weak resistance (**Fig. 4b**) that was in close agreement with previous findings and reflected the well-known preference of the Tat translocase for properly folded substrate proteins ^26, 36^. In stark contrast, the translocase variants each conferred strong resistance (**Fig. 4b**) that was comparable to previously observed suppressor phenotypes.

To corroborate this result, we evaluated the ability of each translocase variant to localize *E. coli* alkaline phosphatase (PhoA) to the periplasm. Previous studies found that PhoA modified with a functional Tat signal peptide (spTorA-PhoA) was only exported by the Tat translocase in mutant *E. coli* strains that permitted oxidative protein folding in the cytoplasm and thus generated correctly folded PhoA moieties prior to export ^26^. Conversely, expression of spTorA-PhoA in the normally reducing cytoplasm of wt *E. coli* resulted in reduced and misfolded PhoA that was blocked for export by wt TatABC. This blockage could be reversed in the presence of Tat QC suppressors that were capable of exporting misfolded spTorA-PhoA out of the reducing cytoplasm of wt *E. coli* ^36^. Here, expression of spTorA-PhoA in the reducing cytoplasm of DADE cells co-expressing H1, H2, or H3 resulted in clearly visible accumulation of PhoA in the periplasm (**Fig. 4c**), corroborating the QC suppressor phenotype for the translocase variants. In contrast, when spTorA-PhoA was expressed in the presence of wt TatABC, there was little detectable PhoA in either the periplasm or the cytoplasm, consistent with previous findings that misfolded PhoA is rejected by wt TatABC and that nonexported PhoA is rapidly degraded ^26, 36^. Taken together, these findings reveal a possible correlation between high-level secretion and QC suppression and suggest that increased translocation flux for H1, H2, and H3 might be gained by relaxation of substrate proofreading.

## Discussion

In this study, we employed a directed co-evolution strategy to isolate gain-of-function Tat translocase variants that enhanced the export of a chimeric reporter protein comprised of a model recombinant antibody, scFv13-R4, fused to Bla. Importantly, three of the isolated variants, H1, H2, and H3, promoted significantly greater secretion of the scFv13-R4-Bla reporter and also its unfused counterpart, scFv13-R4, indicating that the super-secretion phenotype was not limited to the reporter construct. In fact, enhanced export was also observed for several recombinant proteins including antibody-mimetics (*e*.*g*., DARPin, FN3) and a collection of mammalian proteins (*e*.*g*., hGH, hCASP2, hGATA2) and protein domains (*e*.*g*., mEphB2^SAM^, hRAF1^Ras-BD^), demonstrating that the improved secretion capacity of isolated translocase variants was not substrate specific and could be generalized to other structurally diverse proteins that were targeted to the Tat pathway.

Consistent with the low protein flux through the native Tat system ^39, 44, 45^, we determined that the quantity of periplasmic scFv13-R4 that accumulated in cells carrying the H2 translocase was significantly greater than that measured in cells overexpressing wt TatABC. The secreted titer of scFv13-R4 (∼0.5 mg/L) achieved with the H2 translocase compared favorably to the titers obtained using the Sec pathway for periplasmic expression of various scFv proteins ^34, 65^, including Sec-targeted scFv13-R4 ^66^. The titers of hGH purified from the periplasm of cells carrying the H2 translocase were similarly elevated compared to cells carrying wt TatABC, further confirming the universality of the super-secreting H2 translocase. With further optimization of process conditions (*e*.*g*., temperature, inducer concentration, and induction point) and engineering of the Tat translocase in more commonly used industrial strains (*e*.*g*., W3110 and BL21), we expect that the titers of Tat-targeted proteins such as scFv antibodies or hGH could be enhanced beyond the levels reported here or previously for the Sec pathway. In support of this notion is the fact that the Tat system, unlike the Sec pathway, is not essential for cell viability and only handles a relatively small amount of natural protein traffic (∼20-30 substrates are predicted to transit the Tat pathway whereas ∼850 or more proteins are predicted to use the Sec pathway ^23, 24^). In this regard, the Tat system may be more amenable to further expansion of secretion capacity because greater substrate flux would be less likely to compromise cell viability and could proceed without interference from a large numbers of endogenous proteins competing for a limited number of available translocases.

Our efforts to identify mutational alleles that drive high-level secretion through the Tat translocases revealed the importance of cooperativity between the TatB and TatC proteins. This is not entirely surprising given the large number of studies that have reported hetero-oligomeric complexes involving these two proteins. Of relevance here is that TatB and TatC form a functional unit, with the TatBC complex containing an estimated 6–8 copies of each protein and serving as a receptor for signal peptide binding ^23^. Interestingly, the I4V and I14V substitutions acquired by H1 and H2, respectively, occur within the TatB transmembrane helix (TMH) that serves as an important interface with TatC based on previous modelling and experimental studies ^67^. Thus, mutations in these regions may lead to changes at these contact points that alter TatBC complexes in a manner that enhances the translocation rate. In the unique case of H3, our mutational deconvolution revealed that the super-secretion phenotype was attributable to a single P97S substitution in TatC. This residue is located in the first cytoplasmic loop of TatC, a highly conserved region important for recognition of the signal peptide ^62^. Interestingly, translocase variants containing P97S substitutions in TatC were previously isolated as both QC suppressors ^36^, in line with our observation here that H3 exhibited relaxed QC, and as signal peptide suppressors that could export substrates bearing defective signal peptides lacking the twin-arginine motif ^63^.

Interestingly, all three super-secreting translocases acquired greater translocation activity at the expense of the inbuilt QC activity that normally monitors the export readiness of potential substrates by assessing their folding state ^26, 35-38^. This result was somewhat analogous to the trade-off observed in natural and laboratory evolution whereby acquisition of a proficient new enzymatic activity comes at the expense of the original activity or stability ^49^. It is also possible that relaxation of QC is more than just a cost for this new capability, especially if substrate proofreading is a slow step in the translocation process. If that were the case, mutations that reduce or abolish proofreading could lead to faster export kinetics or, conversely, mutations that accelerate export could lead to kinetic bypassing of proofreading whereby substrate export occurs on a much faster timescale than QC. While it remains to be determined which, if any, of these mechanisms underlies super-secretion, such information could make it possible to rationally enhance Tat export even further in the future and could help to unravel the poorly understood phenomenon of how the Tat system synchronizes proofreading of substrate folding with lipid bilayer transport. It also remains to be seen whether enhanced Tat secretion can be achieved with retention of QC activity, which is something that could be investigated by implementing an additional counterselection to the translocase library screening process.

## Materials and Methods

### Bacterial strains and plasmids

*E. coli* strain DH5α was used for all molecular biology while *E. coli* strain DADE (MC4100 Δ*tatABCD*Δ*tatE*) ^51^ lacking all of the *tat* genes was used for all Tat translocation experiments. To evaluate protein export efficiency, this strain was transformed with different pBR322-based plasmids encoding wt or mutant translocases.

### Plasmid construction

All plasmids used in this study are provided in **Supplementary Table 1**. The genetic reporter plasmid pSALect-scFv13-R4 was constructed previously by PCR amplifying the gene encoding scFv13-R4 and cloning it into the MCS of pSALect ^52^ between NdeI and SpeI as described ^33^. A similar approach was used to clone scFv13-R4 into plasmid pDMB ^68^, yielding a plasmid encoding Sec-targeted spDsbA-scFv13-R4-Bla. The plasmids pSALect-off7 and pSALect-YS1 were cloned in a similar manner using the same restriction sites. Plasmid pSALect-scFv-gpD was cloned by PCR amplifying the gene encoding gpD and cloning the resulting product into pSALect between the restriction sites XbaI and SalI. Plasmid pSALect-hGH was cloned by PCR amplifying the gene encoding hGH and cloning the resulting product into pSALect between the restriction sites XbaI and SpeI. Other pSALect-based plasmids used in this study were generated previously ^60^. Plasmid pBC-spTorA-scFv13-R4-FH was generated by PCR amplifying the gene encoding scFv13-R4 with primers that introduced a C-terminal FLAG-6xHis tag and cloning the resulting product between the NdeI and EcoRI restriction sites of pSALect, removing the gene encoding Bla in the process. A similar strategy was used to generate pBC-spDsbA-scFv13-R4-FH. Plasmid pEXT22-spTorA-hGH-6xH was constructed by PCR amplifying the gene encoding spTorA-hGH from pSALect-hGH with primers that introduced a C-terminal 6xHis tag and cloning the resulting product into pEXT22 ^69^ by Gibson assembly. A similar strategy was used to generate pEXT22-spDsbA-hGH-6xH. Plasmid-based expression of TatABC was performed using pTatABC-XX ^36^. In this construct, an XbaI restriction site was introduced between *tatA* and *tatB*, and a XhoI restriction site was introduced between *tatB* and *tatC* by site-directed mutagenesis. These sites were used to generate hybrid translocases where mutant *tatB* or *tatC* alleles were combined with corresponding wt *tat* genes. Individual point mutations of wt TatABC or H3 were introduced using Quik-change site-directed mutagenesis (Agilent) according to manufacturer’s instructions. The same site-directed mutagenesis method was used to mutate the twin-arginine residues in spTorA to twin lysines in pSALect-scFv13-R4.

### Selective plating of bacteria

DADE cells were freshly transformed with one of the pSALect plasmids and one of the pTatABC-XX plasmids carrying either wt or variant translocases and then grown overnight at 37°C in LB containing 20 μg/mL Tc and 30 μg/mL Cm. Cultures were measured to an equivalent Abs_600_ of 1.0 and resuspended in fresh LB media without antibiotics after which cells were serially diluted 10-fold in sterile 96-well plates. A total of 5 μl from the serial dilutions was spotted onto LB-agar containing either Tc and Cm or Carb, after which spots were allowed to dry before plates were incubated at 30°C overnight.

### Selection of translocase variants from combinatorial libraries

To select super-secreting translocase variants, an error-prone library of the *tatABC* operon containing ∼5.5 × 10^6^ members that was generated previously was used here ^36^. Sequencing of a subset of randomly selected clones showed an average mutation rate of two base pairs per gene. The error-prone *tatABC* library was midiprepped and used to transform DADE cells carrying pSALect-scFv13-R4 by electroporation. Transformants were grown in LB with tetracycline (Tc), chloramphenicol (Cm), and 0.2% glucose overnight and serially diluted onto LB-agar plates containing a range of Carb concentrations below and above the minimum inhibitory concentration (MIC) at which cells carrying wt TatABC were confirmed to grow. After incubating the plated library at 30°C overnight, thirty-six colonies that grew under the selection conditions were collected and subjected to back-transformation to ensure that the super-secreting phenotype was associated with the plasmid and not to mutations in the host strain. To back-transform the selected candidates, the pSALect-scFv13-R4 plasmid was cured from the strain by successive growth in the absence of Cm. Once the strains had been cured of pSALect, the plasmids encoding variant translocases were mini-prepped and freshly transformed into DADE cells carrying pSALect-scFv13-R4. Next, cells were spot plated on a range of Carb concentrations and three unique mutants were chosen from among the selected colonies that showed reproducible growth on Carb.

### Subcellular fractionation and Western blotting

For small-scale cultures, cells were grown overnight and subcultured 50-fold in LB containing appropriate antibiotics at 37°C until an absorbance at 600 nm (Abs_600_) of ∼0.5-0.7 was reached, at which time the cultures were induced with 1 mM isopropyl β-d-1-thiogalactopyranoside (IPTG) and incubated at 30°C for an additional 2-4 h. An equivalent amount of cells was harvested by centrifugation at 3,200 rpm for 15 min. Subcellular fractionation was performed by the ice-cold osmotic shock method. Briefly, pelleted cells were resuspended in 1 mL of fractionation buffer consisting of 30 mM Tris-HCl, pH 8.0 1 mM EDTA, and 0.58 M sucrose and left at room temp for 10 min. Samples were centrifuged at 10,000 rpm for 10 min and resuspended in 150 μL of ice-cold 5 mM MgSO_4_ and kept on ice for 10 min. Samples were again centrifuged at 4°C for 10 min at 13,200 rpm. The supernatant was kept as the periplasmic fraction. The remaining pellet was washed once in PBS buffer and resuspended in 300 μL of BugBuster (EMD Millipore) and spun down once more to yield the cytoplasmic fraction from the supernatant. Periplasmic and cytoplasmic fractions were then run on SDS-PAGE gels and Western blotting was performed according to standard protocols using the following antibodies: anti-FLAG-HRP (Abcam ab49763; 1:2,500), anti-PhoA (Millipore MAB1012; 1:2,500), anti-mouse-HRP (Abcam ab6789; 1:5000), anti-GroEL (Sigma G6532; 1:30,000), and anti-rabbit-HRP (Abcam ab205718; 1:5000).

For large-scale cultures, cells were grown overnight and subcultured 50-fold into 1 L of Super Broth (SB) media containing antibiotics at 37°C until an Abs_600_ of ∼0.5-0.7 was reached, at which time cultures were induced with 1 mM IPTG and incubated at 30°C for an additional 16-18 h. An equivalent amount of cells was harvested by centrifugation at 4,500 rpm for 15 min and resuspended in 30 mM Tris-Cl, 0.58 M sucrose, pH 8.0 at 80 mL/g wet weight. Cells were kept on ice while 500 mM EDTA was added dropwise to a final concentration of 1 mM EDTA. Cells were incubated on ice for an additional 10 min with gentle agitation. The cell suspension was centrifuged at 7,000 rpm for 20 min at 4°C and the supernatant was removed. The pellet was resuspended in the same volume of ice-cold 5 mM MgSO_4_ and agitated gently for 10 min in an ice bath. The cell suspension was spun down at 7,000 rpm for 20 min at 4°C, and the supernatant was kept as the osmotic shock fluid containing periplasmic contents. The periplasmic fraction was then used for Ni-NTA purification.

### Protein purification

Osmotic shock fluid containing the periplasmic fraction was incubated with Ni-NTA resin in equilibration buffer (1x PBS buffer with 10 mM imidazole, pH 7.4) for 2 h at 4°C. Proteins were washed and eluted using 1x PBS buffer with 40 mM imidazole and 250 mM imidazole, pH 7.4, respectively. Concentrations were measured by Bradford assay using BSA as standard. Elution fractions were collected and analyzed by SDS-PAGE and subsequent staining with Coomassie blue.

### Enzyme-linked immunosorbent assay (ELISA)

The binding activity of scFv13-R4 was measured by ELISA as described previously ^33^. Briefly, antigen immobilization was performed by treating 96-well EIA plates (Corning) with 100 μL of β-gal (Sigma) at a concentration of 10 μg/mL overnight at 4°C. The plate was washed with PBST (1x PBS, 0.1% Tween20) and blocked with PBST containing 5% (w/v) milk for 2 h at room temp. The plate was washed three times with PBST before incubating with 50 μL of protein that was purified from the periplasmic fractions as described above for 1 h. Plates were washed again three times with PBST and then subjected to anti-FLAG-HRP diluted in PBST (1:5,000) with 1% (w/v) milk for 1 h to determine levels of scFv13-R4 bound to β-gal. Wells were washed a final three times and incubated with Sigma OPD substrate for 30 min. Lastly, wells were quenched with 3N H_2_SO_4_ and the absorbance was read at 492 nm (Abs_492_).

### Cell proliferation assay

The growth stimulatory activity of hGH was determined using an *in vitro* cell proliferation assay as described previously ^70^. Briefly, Ba/F3 cells expressing human growth hormone receptor (Ba/F3-GHR) were starved overnight at 37°C, 5% CO_2_, in RPMI-1640 medium supplemented with 2 mM glutamine, 10% FBS, 2% penicillin-streptomycin and 1 μg/ml puromycin. The starved cells were seeded at a cell density of 1×10^4^ cells/mL in quadruplicates in growth media spiked with equal volumes of purified hGH proteins that had been sterilized by filtration through 0.2 µm membrane filters. Cells were incubated at 37°C, 5% CO_2_ for 72 h, after which the cells density was measured by reading fluorescence at excitation/emission wavelengths of 535/595 nm after being stained by AlmarBlue (Bio-Rad) for 72 h.

## Data availability

All data generated or analyzed during this study are included in this article (and its supplementary information) or are available from the corresponding authors on reasonable request.

## Acknowledgements

We thank Dr. Tracy Palmer for strains and plasmids used in this work. We also thank Dr. Michael Waters and Dr. Andrew Brooks from The University of Queensland, Australia for providing the Ba/F3-GHR cell line used in this study. This work was supported by the National Science Foundation (grants # CBET-0449080 and CBET-1605242 to M.P.D.), the National Institutes of Health (grant # CA132223A (to M.P.D.), the New York State Office of Science, Technology and Academic Research Distinguished Faculty Award (to M.P.D.), and a Royal Thai Government Fellowship (to D.W.-Z.).

## Author Contributions

M.N.T. designed research, performed research, analyzed data, and wrote the paper. M.L., D.K., M.A.R., and D.W-Z. designed research, performed research, and analyzed data. M.P.D. designed and directed research, analyzed data, and wrote the paper.

## References

1. Georgiou, G., Segatori, L., Preparative expression of secreted proteins in bacteria: status report and future prospects. Curr Opin Biotechnol 2005, 16 (5), 538–45.

2. Baneyx, F., Mujacic, M., Recombinant protein folding and misfolding in Escherichia coli. Nat Biotechnol 2004, 22 (11), 1399–408.

3. Choi, J. H., Lee, S. Y., Secretory and extracellular production of recombinant proteins using Escherichia coli. Appl Microbiol Biotechnol 2004, 64 (5), 625–35.

4. Smith, G. P., Filamentous fusion phage: novel expression vectors that display cloned antigens on the virion surface. Science 1985, 228 (4705), 1315–7.

5. Francisco, J. A., Campbell, R., Iverson, B. L., Georgiou, G., Production and fluorescence-activated cell sorting of Escherichia coli expressing a functional antibody fragment on the external surface. Proc Natl Acad Sci U S A 1993, 90 (22), 10444–8.

6. Chen, G., Hayhurst, A., Thomas, J. G., Harvey, B. R., Iverson, B. L., Georgiou, G., Isolation of high-affinity ligand-binding proteins by periplasmic expression with cytometric screening (PECS). Nat Biotechnol 2001, 19 (6), 537–42.

7. Mazor, Y., Van Blarcom, T., Mabry, R., Iverson, B. L., Georgiou, G., Isolation of engineered, full-length antibodies from libraries expressed in Escherichia coli. Nat Biotechnol 2007, 25 (5), 563–5.

8. Fisher, A. C., Haitjema, C. H., Guarino, C., Celik, E., Endicott, C. E., Reading, C. A., Merritt, J. H., Ptak, A. C., Zhang, S., DeLisa, M. P., Production of secretory and extracellular N-linked glycoproteins in Escherichia coli. Appl Environ Microbiol 2011, 77 (3), 871–81.

9. Valderrama-Rincon, J. D., Fisher, A. C., Merritt, J. H., Fan, Y. Y., Reading, C. A., Chhiba, K., Heiss, C., Azadi, P., Aebi, M., DeLisa, M. P., An engineered eukaryotic protein glycosylation pathway in Escherichia coli. Nat Chem Biol 2012, 8 (5), 434–6.

10. Wacker, M., Linton, D., Hitchen, P. G., Nita-Lazar, M., Haslam, S. M., North, S. J., Panico, M., Morris, H. R., Dell, A., Wren, B. W., Aebi, M., N-linked glycosylation in Campylobacter jejuni and its functional transfer into E. coli. Science 2002, 298 (5599), 1790–3.

11. Fisher, A. C., Kim, W., DeLisa, M. P., Genetic selection for protein solubility enabled by the folding quality control feature of the twin-arginine translocation pathway. Protein Sci 2006, 15 (3), 449–58.

12. Karlsson, A. J., Lim, H. K., Xu, H., Rocco, M. A., Bratkowski, M. A., Ke, A., DeLisa, M. P., Engineering antibody fitness and function using membrane-anchored display of correctly folded proteins. J Mol Biol 2012, 416 (1), 94–107.

13. Waraho, D., DeLisa, M. P., Versatile selection technology for intracellular protein-protein interactions mediated by a unique bacterial hitchhiker transport mechanism. Proc Natl Acad Sci U S A 2009, 106 (10), 3692–7.

14. Schatz, G., Dobberstein, B., Common principles of protein translocation across membranes. Science 1996, 271 (5255), 1519–26.

15. Driessen, A. J., Nouwen, N., Protein translocation across the bacterial cytoplasmic membrane. Annu Rev Biochem 2008, 77, 643–67.

16. Benson, S. A., Bremer, E., Silhavy, T. J., Intragenic regions required for LamB export. Proc Natl Acad Sci U S A 1984, 81 (12), 3830–4.

17. Kiino, D. R., Silhavy, T. J., Mutation prlF1 relieves the lethality associated with export of beta-galactosidase hybrid proteins in Escherichia coli. J Bacteriol 1984, 158 (3), 878–83.

18. Hayhurst, A., Georgiou, G., High-throughput antibody isolation. Curr Opin Chem Biol 2001, 5 (6), 683–9.

19. Schierle, C. F., Berkmen, M., Huber, D., Kumamoto, C., Boyd, D., Beckwith, J., The DsbA signal sequence directs efficient, cotranslational export of passenger proteins to the Escherichia coli periplasm via the signal recognition particle pathway. J Bacteriol 2003, 185 (19), 5706–13.

20. Gentz, R., Kuys, Y., Zwieb, C., Taatjes, D., Taatjes, H., Bannwarth, W., Stueber, D., Ibrahimi, I., Association of degradation and secretion of three chimeric polypeptides in Escherichia coli. J Bacteriol 1988, 170 (5), 2212–20.

21. Feilmeier, B. J., Iseminger, G., Schroeder, D., Webber, H., Phillips, G. J., Green fluorescent protein functions as a reporter for protein localization in Escherichia coli. J Bacteriol 2000, 182 (14), 4068–76.

22. Natale, P., Bruser, T., Driessen, A. J., Sec-and Tat-mediated protein secretion across the bacterial cytoplasmic membrane--distinct translocases and mechanisms. Biochim Biophys Acta 2008, 1778 (9), 1735–56.

23. Palmer, T., Berks, B. C., The twin-arginine translocation (Tat) protein export pathway. Nat Rev Microbiol 2012, 10 (7), 483–96.

24. Berks, B. C., Sargent, F., Palmer, T., The Tat protein export pathway. Mol Microbiol 2000, 35 (2), 260–74.

25. Frain, K. M., van Dijl, J. M., Robinson, C., The twin-arginine pathway for protein secretion. EcoSal Plus 2019, 8 (2).

26. DeLisa, M. P., Tullman, D., Georgiou, G., Folding quality control in the export of proteins by the bacterial twin-arginine translocation pathway. Proc Natl Acad Sci U S A 2003, 100 (10), 6115–20.

27. Thomas, J. D., Daniel, R. A., Errington, J., Robinson, C., Export of active green fluorescent protein to the periplasm by the twin-arginine translocase (Tat) pathway in Escherichia coli. Mol Microbiol 2001, 39 (1), 47–53.

28. Lee, H. C., Portnoff, A. D., Rocco, M. A., DeLisa, M. P., An engineered genetic selection for ternary protein complexes inspired by a natural three-component hitchhiker mechanism. Sci Rep 2014, 4, 7570.

29. Rodrigue, A., Chanal, A., Beck, K., Muller, M., Wu, L. F., Co-translocation of a periplasmic enzyme complex by a hitchhiker mechanism through the bacterial tat pathway. J Biol Chem 1999, 274 (19), 13223–8.

30. Kim, J.-Y., Fogarty, E. A., Lu, F. J., Zhu, H., Henderson, L. A., DeLisa, M. P., Twin-arginine translocation of active human tissue plasminogen activator in Escherichia coli. Appl Environ Microbiol 2005, 71, 8451–8459.

31. Matos, C. F., Robinson, C., Alanen, H. I., Prus, P., Uchida, Y., Ruddock, L. W., Freedman, R. B., Keshavarz-Moore, E., Efficient export of prefolded, disulfide-bonded recombinant proteins to the periplasm by the Tat pathway in Escherichia coli CyDisCo strains. Biotechnol Prog 2014, 30 (2), 281–90.

32. Alanen, H. I., Walker, K. L., Lourdes Velez Suberbie, M., Matos, C. F., Bonisch, S., Freedman, R. B., Keshavarz-Moore, E., Ruddock, L. W., Robinson, C., Efficient export of human growth hormone, interferon alpha2b and antibody fragments to the periplasm by the Escherichia coli Tat pathway in the absence of prior disulfide bond formation. Biochim Biophys Acta 2015, 1853 (3), 756–63.

33. Fisher, A. C., DeLisa, M. P., Efficient isolation of soluble intracellular single-chain antibodies using the twin-arginine translocation machinery. J Mol Biol 2009, 385 (1), 299–311.

34. Ribnicky, B., Van Blarcom, T., Georgiou, G., A scFv antibody mutant isolated in a genetic screen for improved export via the twin arginine transporter pathway exhibits faster folding. J Mol Biol 2007, 369 (3), 631–9.

35. Jones, A. S., Austerberry, J. I., Dajani, R., Warwicker, J., Curtis, R., Derrick, J. P., Robinson, C., Proofreading of substrate structure by the twin-arginine translocase is highly dependent on substrate conformational flexibility but surprisingly tolerant of surface charge and hydrophobicity changes. Biochim Biophys Acta 2016, 1863 (12), 3116–3124.

36. Rocco, M. A., Waraho-Zhmayev, D., DeLisa, M. P., Twin-arginine translocase mutations that suppress folding quality control and permit export of misfolded substrate proteins. Proc Natl Acad Sci U S A 2012, 109 (33), 13392–7.

37. Sutherland, G. A., Grayson, K. J., Adams, N. B. P., Mermans, D. M. J., Jones, A. S., Robertson, A. J., Auman, D. B., Brindley, A. A., Sterpone, F., Tuffery, P., Derreumaux, P., Dutton, P. L., Robinson, C., Hitchcock, A., Hunter, C. N., Probing the quality control mechanism of the Escherichia coli twin-arginine translocase with folding variants of a de novo-designed heme protein. J Biol Chem 2018, 293 (18), 6672–6681.

38. Taw, M. N., Boock, J. T., Kim, D., Rocco, M. A., Waraho-Zhmayev, D., DeLisa, M. P., Twin-arginine translocase component TatB performs folding quality control via a general chaperone activity. bioRxiv 2020, 2020.05.11.089458.

39. Santini, C. L., Ize, B., Chanal, A., Muller, M., Giordano, G., Wu, L. F., A novel sec-independent periplasmic protein translocation pathway in Escherichia coli. EMBO J 1998, 17 (1), 101–12.

40. Sanders, C., Wethkamp, N., Lill, H., Transport of cytochrome c derivatives by the bacterial Tat protein translocation system. Mol Microbiol 2001, 41 (1), 241–6.

41. Bruser, T., Yano, T., Brune, D. C., Daldal, F., Membrane targeting of a folded and cofactor-containing protein. Eur J Biochem 2003, 270 (6), 1211–21.

42. Maurer, C., Panahandeh, S., Moser, M., Muller, M., Impairment of twin-arginine-dependent export by seemingly small alterations of substrate conformation. FEBS Lett 2009, 583 (17), 2849–53.

43. Fisher, A. C., Kim, J.-Y., Perez-Rodriguez, R., Tullman-Ercek, D., Fish, W., Henderson, L. A., DeLisa, M. P., Exploration of twin-arginine translocation for the expression and purification of correctly folded proteins in Escherichia coli. Microbial Biotechnol 2008, 1, 403–15.

44. Sargent, F., Bogsch, E. G., Stanley, N. R., Wexler, M., Robinson, C., Berks, B. C., Palmer, T., Overlapping functions of components of a bacterial Sec-independent protein export pathway. EMBO J 1998, 17 (13), 3640–50.

45. Bogsch, E. G., Sargent, F., Stanley, N. R., Berks, B. C., Robinson, C., Palmer, T., An essential component of a novel bacterial protein export system with homologues in plastids and mitochondria. J Biol Chem 1998, 273 (29), 18003–6.

46. Matos, C. F., Branston, S. D., Albiniak, A., Dhanoya, A., Freedman, R. B., Keshavarz-Moore, E., Robinson, C., High-yield export of a native heterologous protein to the periplasm by the tat translocation pathway in Escherichia coli. Biotechnol Bioeng 2012, 109 (10), 2533–42.

47. Browning, D. F., Richards, K. L., Peswani, A. R., Roobol, J., Busby, S. J. W., Robinson, C., Escherichia coli “TatExpress” strains super-secrete human growth hormone into the bacterial periplasm by the Tat pathway. Biotechnol Bioeng 2017, 114 (12), 2828–2836.

48. Guerrero Montero, I., Richards, K. L., Jawara, C., Browning, D. F., Peswani, A. R., Labrit, M., Allen, M., Aubry, C., Dave, E., Humphreys, D. P., Busby, S. J. W., Robinson, C., Escherichia coli “TatExpress” strains export several g/L human growth hormone to the periplasm by the Tat pathway. Biotechnol Bioeng 2019, 116 (12), 3282–3291.

49. Khersonsky, O., Tawfik, D. S., Enzyme promiscuity: a mechanistic and evolutionary perspective. Annu Rev Biochem 2010, 79, 471–505.

50. Martineau, P., Jones, P., Winter, G., Expression of an antibody fragment at high levels in the bacterial cytoplasm. J Mol Biol 1998, 280 (1), 117–27.

51. Wexler, M., Sargent, F., Jack, R. L., Stanley, N. R., Bogsch, E. G., Robinson, C., Berks, B. C., Palmer, T., TatD is a cytoplasmic protein with DNase activity. No requirement for TatD family proteins in sec-independent protein export. J Biol Chem 2000, 275 (22), 16717–22.

52. Lutz, S., Fast, W., Benkovic, S. J., A universal, vector-based system for nucleic acid reading-frame selection. Protein Eng 2002, 15 (12), 1025–30.

53. Zhang, Y., Wang, L., Hu, Y., Jin, C., Solution structure of the TatB component of the twin-arginine translocation system. Biochim Biophys Acta 2014, 1838 (7), 1881–8.

54. Rollauer, S. E., Tarry, M. J., Graham, J. E., Jaaskelainen, M., Jager, F., Johnson, S., Krehenbrink, M., Liu, S. M., Lukey, M. J., Marcoux, J., McDowell, M. A., Rodriguez, F., Roversi, P., Stansfeld, P. J., Robinson, C. V., Sansom, M. S., Palmer, T., Hogbom, M., Berks, B. C., Lea, S. M., Structure of the TatC core of the twin-arginine protein transport system. Nature 2012, 492 (7428), 210–4.

55. Ize, B., Stanley, N. R., Buchanan, G., Palmer, T., Role of the Escherichia coli Tat pathway in outer membrane integrity. Mol Microbiol 2003, 48 (5), 1183–93.

56. Koch, H., Grafe, N., Schiess, R., Pluckthun, A., Direct selection of antibodies from complex libraries with the protein fragment complementation assay. J Mol Biol 2006, 357 (2), 427–41.

57. Binz, H. K., Amstutz, P., Kohl, A., Stumpp, M. T., Briand, C., Forrer, P., Grutter, M. G., Pluckthun, A., High-affinity binders selected from designed ankyrin repeat protein libraries. Nat Biotechnol 2004, 22 (5), 575–82.

58. Koide, A., Bailey, C. W., Huang, X., Koide, S., The fibronectin type III domain as a scaffold for novel binding proteins. J Mol Biol 1998, 284 (4), 1141–51.

59. Hsiung, H. M., Becker, G. W., Secretion and folding of human growth hormone in Escherichia coli. Biotechnol Genet Eng Rev 1988, 6, 43–65.

60. Lim, H. K., Mansell, T. J., Linderman, S. W., Fisher, A. C., Dyson, M. R., DeLisa, M. P., Mining mammalian genomes for folding competent proteins using Tat-dependent genetic selection in Escherichia coli. Protein Sci 2009, 18 (12), 2537–49.

61. Youngman, K. M., Spencer, D. B., Brems, D. N., DeFelippis, M. R., Kinetic analysis of the folding of human growth hormone. Influence of disulfide bonds. J Biol Chem 1995, 270 (34), 19816–22.

62. Zoufaly, S., Frobel, J., Rose, P., Flecken, T., Maurer, C., Moser, M., Muller, M., Mapping precursor-binding site on TatC subunit of twin arginine-specific protein translocase by site-specific photo cross-linking. J Biol Chem 2012, 287 (16), 13430–41.

63. Strauch, E. M., Georgiou, G., Escherichia coli tatC mutations that suppress defective twin-arginine transporter signal peptides. J Mol Biol 2007, 374 (2), 283–91.

64. Bryson, J. W., Desjarlais, J. R., Handel, T. M., DeGrado, W. F., From coiled coils to small globular proteins: design of a native-like three-helix bundle. Protein Sci 1998, 7 (6), 1404–14.

65. Kipriyanov, S. M., Moldenhauer, G., Little, M., High level production of soluble single chain antibodies in small-scale Escherichia coli cultures. J Immunol Methods 1997, 200 (1-2), 69–77.

66. Kasli, I. M., Thomas, O. R. T., Overton, T. W., Use of a design of experiments approach to optimise production of a recombinant antibody fragment in the periplasm of Escherichia coli: selection of signal peptide and optimal growth conditions. AMB Express 2019, 9 (1), 5.

67. Alcock, F., Stansfeld, P. J., Basit, H., Habersetzer, J., Baker, M. A., Palmer, T., Wallace, M. I., Berks, B. C., Assembling the Tat protein translocase. Elife 2016, 5.

68. Mansell, T. J., Linderman, S. W., Fisher, A. C., DeLisa, M. P., A rapid protein folding assay for the bacterial periplasm. Protein Sci 2010, 19 (5), 1079–90.

69. Dykxhoorn, D. M., St Pierre, R., Linn, T., A set of compatible tac promoter expression vectors. Gene 1996, 177 (1-2), 133–6.

70. Behncken, S. N., Rowlinson, S. W., Rowland, J. E., Conway-Campbell, B. L., Monks, T. A., Waters, M. J., Aspartate 171 is the major primate-specific determinant of human growth hormone. Engineering porcine growth hormone to activate the human receptor. J Biol Chem 1997, 272 (43), 27077–83.

