## Supplemental Table and Figures for "Engineering a super-secreting strain of *Escherichia coli* by directed co-evolution of the multiprotein Tat translocation machinery"

**Supplementary Table 1. Bacterial strains and plasmids used in this study**

| Bacterial strain | Genotype | Source |
|---|---|---|
| MC4100 | $F^-$ *araD139* Δ(*argF-lac*)*169* $λ^-$ *e14*$^-$ *flhD5301* Δ(*fruK-yeiR*)*725(fruA25)* *relA1 rpsL150*(Str$^R$) *rbsR22* Δ(*fimB*-fimE)632(::IS*1*) *deoC1* | 1 |
| DADE | MC4100 Δ*tatABCD*Δ*tatE* | 2 |
| **Bacterial plasmids** | | |
| pTatABC | Entire *E. coli tatABC* operon including native promoter in pBR322; Tet$^R$ | 3 |
| pTatABC-XX | pTatABC with XbaI and XhoI restriction sites flanking *tatB* and *tatC* genes, respectively | 3 |
| pTatABC-H1 | pTatABC-XX but with genes encoding H1 translocase variant | This study |
| pTatABC-H2 | pTatABC-XX but with genes encoding H2 translocase variant | This study |
| pTatABC-H3 | pTatABC-XX but with genes encoding H3 translocase variant | This study |
| pSALect-scFv13-R4 | Gene encoding scFv13-R4 protein cloned between spTorA signal peptide and TEM1-Bla reporter in plasmid pSALect, a pBC-SK(+) derivative; *lac* promoter, Cam$^R$, Amp$^R$ | 4 |
| pSALect-KK-scFv13-R4 | Same as pSALect-scFv13-R4 but with twin arginine residues in spTorA signal peptide mutated to twin lysines | This study |
| pDMB-scFv13-R4 | Gene encoding scFv13-R4 protein cloned between spDsbA signal peptide and TEM1-Bla reporter in plasmid pDMB; Cam$^R$, Amp$^R$ | This study |
| pSALect-gpD | Gene encoding bacteriophage lambda gpD protein cloned between spTorA and TEM1-Bla in plasmid pSALect | This study |
| pSALect-off7 | Gene encoding DARPin off7 protein cloned between spTorA and TEM1-Bla in plasmid pSALect | This study |
| pSALect-YS1 | Gene encoding FN3 monobody YS1 protein cloned between spTorA and TEM1-Bla in plasmid pSALect | This study |
| pSALect-hGH | Gene encoding hGH cloned between spTorA and TEM1-Bla in plasmid pSALect | This study |
| pSALect-hCASP2 | Gene encoding human CASP2 protease cloned between spTorA and TEM1-Bla in plasmid pSALect | 5 |
| pSALect-mEphB2$^{SAM}$ | Gene encoding sterile alpha motif (SAM) domain of murine EphB2 protein cloned between spTorA and TEM1-Bla in plasmid pSALect | 5 |
| pSALect-hGATA2 | Gene encoding human GATA2 transcription factor cloned between spTorA and TEM1-Bla in plasmid pSALect | 5 |
| pSALect-hRAF1$^{Ras\text{-}BD}$ | Gene encoding Ras binding domain (Ras-BD) of human RAF1 protein cloned between spTorA and TEM1-Bla in plasmid pSALect | 5 |
| pSALect-α3A | Codon-optimized gene encoding α3A protein cloned between spTorA and TEM1-Bla in plasmid pSALect | 3 |
| pBC-spTorA-scFv13-R4-FH | Gene encoding scFv13-R4 protein modified with N-terminal spTorA signal peptide and C-terminal FLAG-6xHis in plasmid pBC-SK(+) | This study |
| pBC-spDsbA-scFv13-R4-FH | Same as pBC-spTorA-scFv13-R4 but with N-terminal spDsbA signal peptide | This study |
| pEXT22-spTorA-hGH-6xHis | Gene encoding hGH modified with N-terminal spTorA signal peptide and C-terminal 6xHis in plasmid pEXT22; *tac* promoter, Kan$^R$ | This study |
| pEXT22-spDsbA-hGH-6xHis | Same as pEXT22-spTorA-hGH but with N-terminal spDsbA signal peptide | This study |
| pTorA-AP | Signal peptide derived from *E. coli* trimethylamine *N*-oxide reductase (TorA) fused to gene encoding mature *E. coli* alkaline phosphatase (APΔ1-22) cloned in plasmid pTrc99A | 6 |
| pSALect-α3 plasmids | Codon-optimized genes encoding α3A/B/C/D proteins cloned between spTorA and Bla in plasmid pSALect | 3 |
| pTatAB$^{H1}$C | pTatABC-XX but with gene encoding *tatB* allele of H1 translocase variant | This study |
| pTatAB$^{H2}$C | pTatABC-XX but with gene encoding *tatB* allele of H2 translocase variant | This study |
| pTatABC$^{H1}$ | pTatABC-XX but with gene encoding *tatC* allele of H1 translocase variant | This study |
| pTatABC$^{H2}$ | pTatABC-XX but with gene encoding *tatC* allele of H2 translocase variant | This study |
| pTatABC$^{P97S}$ | pTatABC-XX but with gene encoding *tatC* with P97S substitution | This study |
| pTatABC-H3(TatC$^{T14I}$) | pTatABC-H3 but with T14I reversion mutation in *tatC* | This study |

| | | |
|---|---|---|
| pTatABC-H3(TatC$^{A188D}$) | pTatABC-H3 but with A188D reversion mutation in *tatC* | This study |
| pTatABC-H3(TatC$^{A202V}$) | pTatABC-H3 but with A202V reversion mutation in *tatC* | This study |
| pTatABC-H3(TatC$^{L243R}$) | pTatABC-H3 but with L243R reversion mutation in *tatC* | This study |
| pTatABC-H3(TatC$^{S97P}$) | pTatABC-H3 but with S97P reversion mutation in *tatC* | This study |

**References cited**

1. Bogsch, E. G.; Sargent, F.; Stanley, N. R.; Berks, B. C.; Robinson, C.; Palmer, T., An essential component of a novel bacterial protein export system with homologues in plastids and mitochondria. *J Biol Chem* **1998,** *273* (29), 18003-6.
2. Wexler, M.; Sargent, F.; Jack, R. L.; Stanley, N. R.; Bogsch, E. G.; Robinson, C.; Berks, B. C.; Palmer, T., TatD is a cytoplasmic protein with DNase activity - No requirement for TatD family proteins in Sec-independent protein export. *J Biol Chem* **2000,** *275* (22), 16717-16722.
3. Rocco, M. A.; Waraho-Zhmayev, D.; DeLisa, M. P., Twin-arginine translocase mutations that suppress folding quality control and permit export of misfolded substrate proteins. *Proc Natl Acad Sci U S A* **2012,** *109* (33), 13392-13397.
4. Fisher, A. C.; DeLisa, M. P., Efficient isolation of soluble intracellular single-chain antibodies using the twin-arginine translocation machinery. *J Mol Biol* **2009,** *385* (1), 299-311.
5. Lim, H. K.; Mansell, T. J.; Linderman, S. W.; Fisher, A. C.; Dyson, M. R.; DeLisa, M. P., Mining mammalian genomes for folding competent proteins using Tat-dependent genetic selection in Escherichia coli. *Protein Sci* **2009,** *18* (12), 2537-49.
6. DeLisa, M. P.; Tullman, D.; Georgiou, G., Folding quality control in the export of proteins by the bacterial twin-arginine translocation pathway. *Proc Natl Acad Sci U S A* **2003,** *100* (10), 6115-6120.

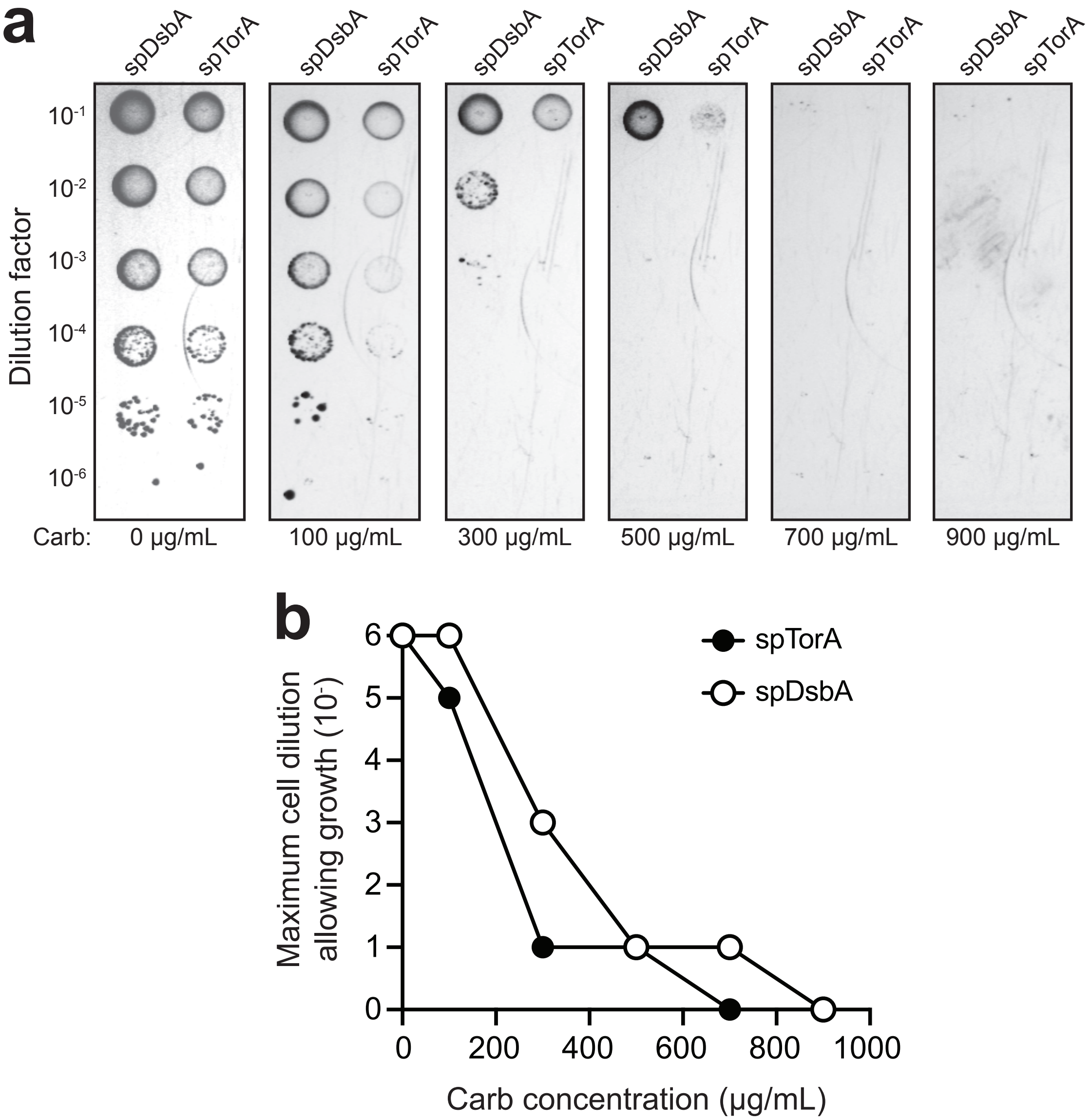


**Supplementary Figure 1. Tat- versus Sec-mediated export of scFv13-R4-Bla reporter.** (a) Spot plating and (b) corresponding kill curves corresponding to DADE cells co-expressing either spDsbA-scFv13-R4-Bla or spTorA-scFv13-R4-Bla along with plasmid-encoded copies of wt TatABC. Serially diluted cells were spotted on LB-agar plates supplemented with varying concentrations of carbenicillin (Carb; 100-900 µg/mL) or 20 µg/mL tetracycline and 30 µg/mL chloramphenicol (0 µg/mL Carb).

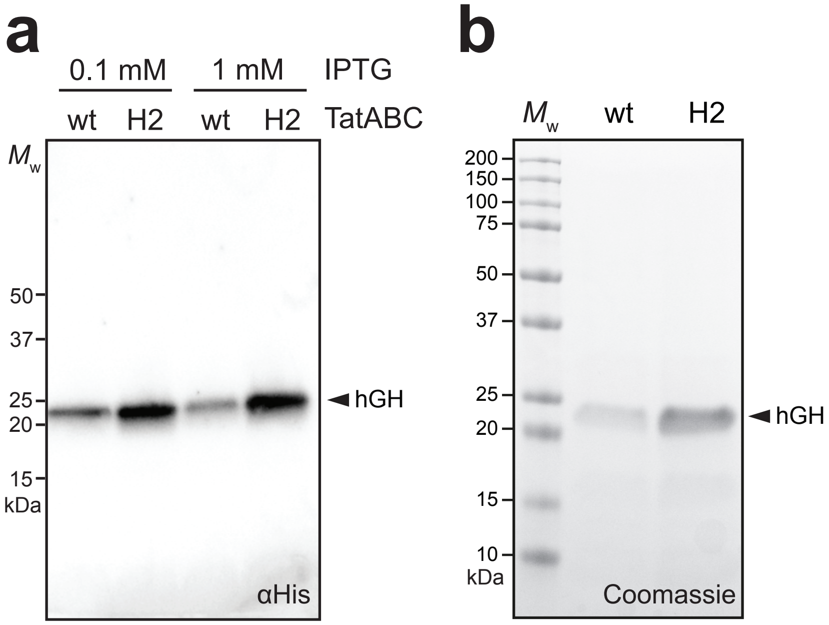


**Supplementary Figure 2. Expression and purification of hGH by H2 translocase.** (a) Western blot analysis of periplasmic fractions prepared from DADE cells co-expressing the 6xHis-tagged version of spTorA-hGH along with either wt TatABC or the H2 translocase mutant as indicated. Fractions corresponding to an equivalent number of cells were loaded in each lane. Anti-hexahistidine (αHis) antibody was used to detect hGH. Molecular weight (MW) marker is indicated on the left. (b) Coomassie-stained SDS-PAGE gel showing spTorA-hGH purified from the periplasm of DADE cells co-expressing either wt or H2 translocases as indicated. Molecular weight (MW) marker included on the left. Protein titers were calculated by densitometry analysis of protein bands corresponding to hGH using ChemiDoc XRS+ System with ImageLab software. Images in (a) and (b) are representative of three biological replicates.

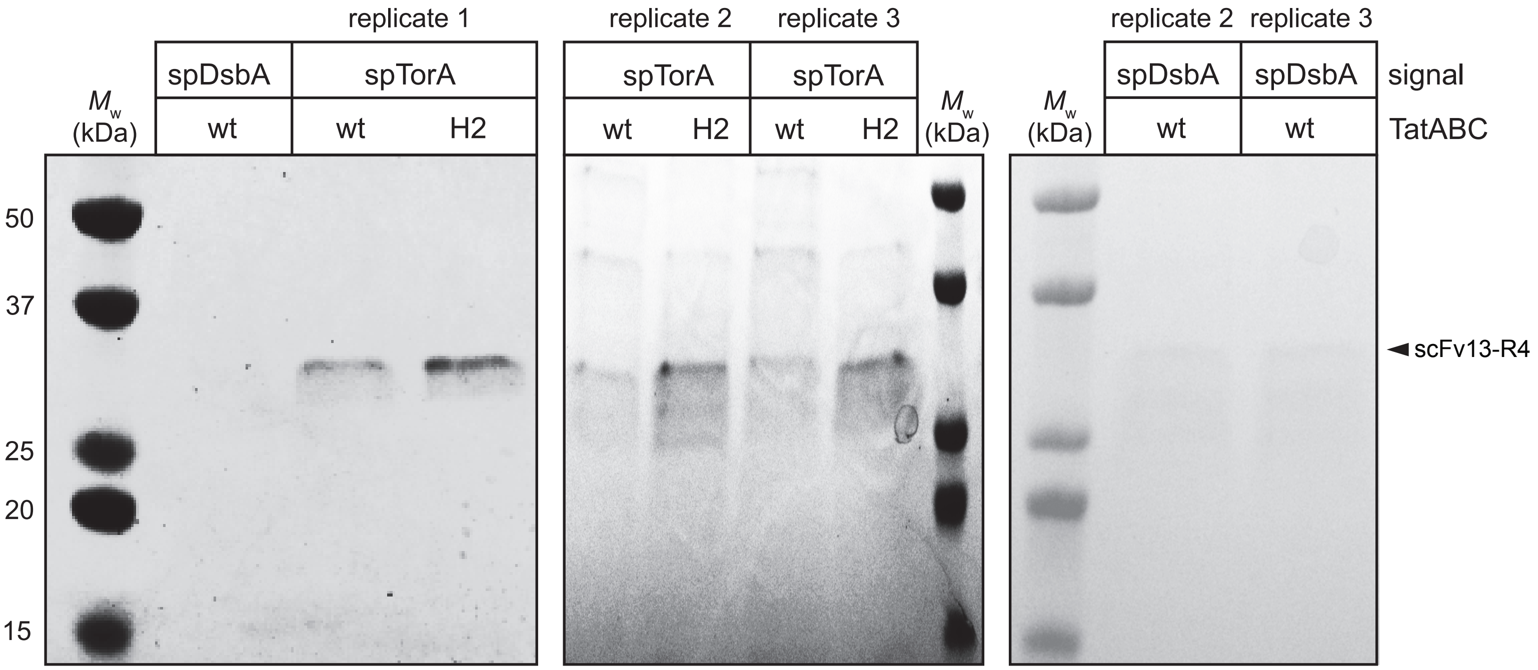


**Supplementary Figure 3. Purification of scFv13-R4 following export by H2 translocase.** Coomassie-stained SDS-PAGE gels showing spDsbA-scFv13-R4 or spTorA-scFv13-R4 proteins purified from the periplasm of DADE cells co-expressing either wt TatABC or the H2 translocase mutant as indicated. Molecular weight (MW) marker included on the left. SDS-PAGE gels depict three biological replicates. Protein titers were calculated by densitometry analysis of protein bands corresponding to scFv13-R4 using ChemiDoc XRS+ System with ImageLab software.

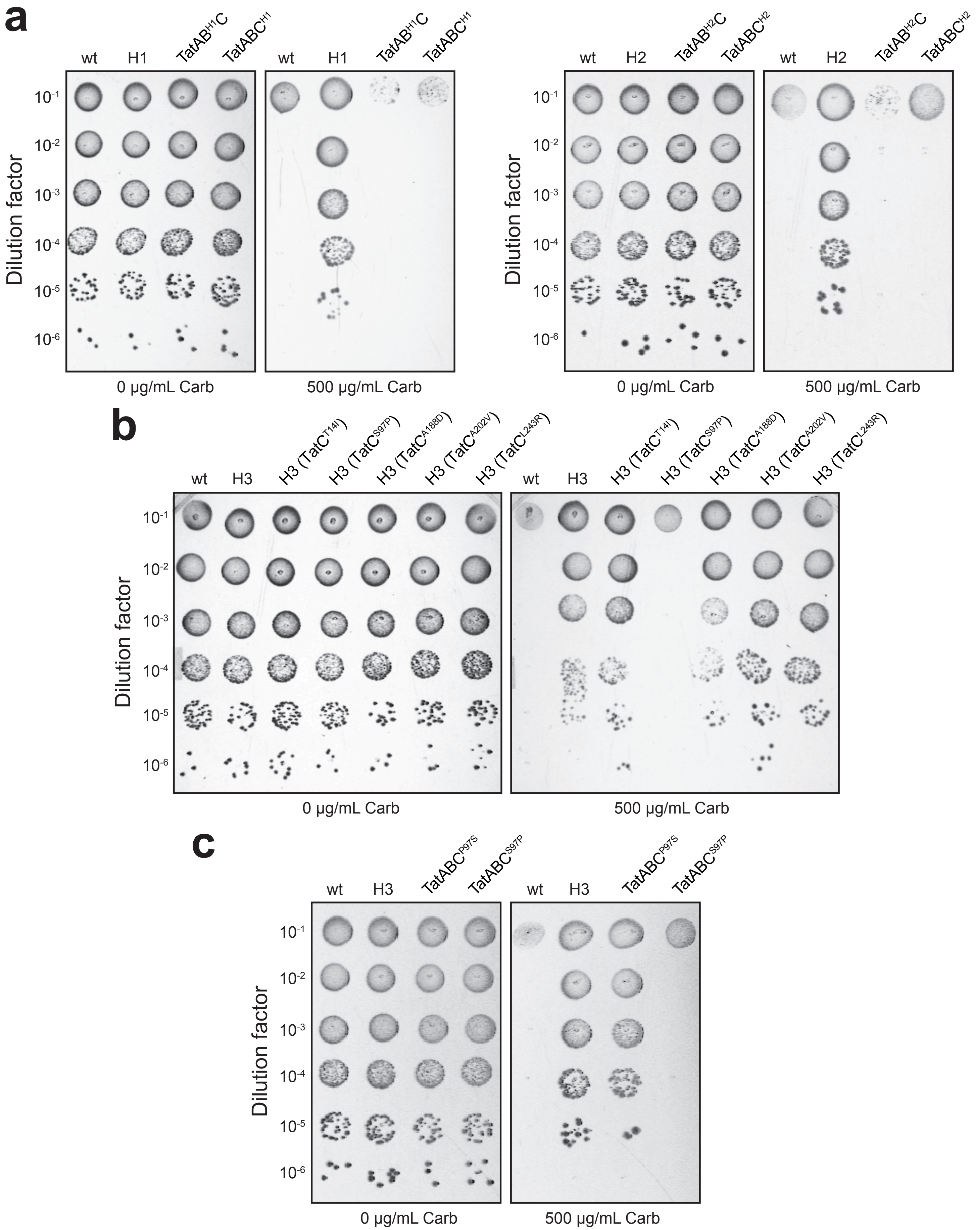


**Supplementary Figure 4. Resistance conferred by hybrid translocase variants.** (a-c) Spot plating of DADE cells co-expressing spTorA-scFv13-R4-Bla along with plasmid-encoded copies of hybrid translocases created by replacing one of the mutated *tat* alleles from each mutant with the corresponding wt alleles. For example, combining mutant *tatB* allele from H1 with wt *tatAC* alleles gave rise to TatAB$^{H1}$C translocase. Cells co-expressing wt TatABC or H1/H2/H3 translocases were included as controls. Serially diluted cells were spotted on LB-agar plates supplemented with carbenicillin (500 µg/mL Carb) or 20 µg/mL tetracycline and 30 µg/mL chloramphenicol (0 µg/mL Carb).
